# Polymer physics-based classification of neurons

**DOI:** 10.1101/2022.04.07.487455

**Authors:** Kiri Choi, Won Kyu Kim, Changbong Hyeon

## Abstract

Recognizing that diverse morphologies of neurons are reminiscent of structures of branched polymers, we put forward a principled and systematic way of classifying neurons that employs the ideas of polymer physics. In particular, we use 3D coordinates of individual neurons, which are accessible in recent neuron reconstruction datasets from electron microscope images. We numerically calculate the form factor, *F* (*q*), a Fourier transform of the distance distribution of particles comprising an object of interest, which is routinely measured in scattering experiments to quantitatively characterize the structure of materials. For a polymer-like object consisting of *n* monomers spanning over a length scale of *r, F* (*q*) scales with the wavenumber *q*(= 2*π/r*) as *F* (*q*) ∼ *q*^−𝒟^ at an intermediate range of *q*, where 𝒟 is the fractal dimension or the inverse scaling exponent (𝒟 = *ν*^−1^) characterizing the geometrical feature (*r* ∼ *n*^*ν*^) of the object. *F* (*q*) can be used to describe a neuron morphology in terms of its size (*R*_*n*_) and the extent of branching quantified by 𝒟. By defining the distance between *F* (*q*)s as a measure of similarity between two neuronal morphologies, we tackle the neuron classification problem. In comparison with other existing classification methods for neuronal morphologies, our *F* (*q*)-based classification rests solely on 3D coordinates of neurons with no prior knowledge of morphological features. When applied to publicly available neuron datasets from three different organisms, our method not only complements other methods but also offers a physical picture of how the dendritic and axonal branches of an individual neuron fill the space of dense neural networks inside the brain.

## INTRODUCTION

Form determines function in biology. Neurons, which are the basic signaling units of the nervous system, are not an exception [1]. Since Ramón y Cajal provided evidence that this general hypothesis was also at work in neurons [2], the problem of neuron classification based on their morphology has been a subject of considerable interest in brain research. Along with a causal relationship for the correlation between the neuronal morphologies and spiking patterns of electrophysiological recordings [3], the morphological detail of a neuron has been suggested as one of the key determinants of physiology and functional differentiation of neurons [4–8], engendering a number of morphology-based classification methods [9–23].

Neurons, demonstrating a variety of elaborate arborization and branching patterns, are considered fractal objects, which has led several studies to calculate fractal dimensions to characterize neurons by the box-counting method [7, 24–28]. As a physical object, many neurons entangled and filling the space inside the brain are reminiscent of a solution or a melt of branched polymers whose space-filling structure can also be quantitatively mapped to the corresponding fractal dimension. Scattering experiments have long been employed to study the structures of polymer chains in various solvent conditions or polymer concentrations [29, 30], and they can also offer the fractal dimension of polymer chains. Thus, it would be natural to analyze the neuronal morphologies by utilizing the ideas of scattering experiments. To be specific, we put forward calculating the form factor, *F* (*q*), to quantify the structures of individual neurons, calculate the fractal dimension of neurons just like the analyses done for polymer chains, and then grouping the neurons based on the similarity between the calculated *F* (*q*)s.

Here we perform our *F* (*q*)-based analysis on publicly available datasets of neuron morphologies obtained for (i) *C. elegans* nervous system [31], (ii) *Drosophila* olfactory projection neurons [13], and (iii) the mouse primary visual cortex (V1) neurons in the Allen Cell Type database [11]. The *F* (*q*)based analysis leverages the full 3D coordinates of neurons reconstructed from electron microscope images without resorting to any prior knowledge of the neuron morphology. The outcomes from our classification method are found comparable to other existing analyses; yet, they offer novel insights into neuron morphologies in the language of branched polymers under various conditions.

### THEORETICAL BACKGROUND

Here we overview basic concepts of polymer physics and key quantities, such as scaling exponent (*ν*) and fractal dimension (𝒟 = 1*/ν*), that we will use to describe the neuron morphologies. A description of two length scales in neuron morphology is given in the first subsection. The second subsection provides the definition of the form factor. The remaining two subsections describe the basic argument by Flory that is employed to determine the scaling relationship of polymer size (Flory radius, *R*_ℱ_) with the polymer length (*N*), i.e., *R*_ℱ_ ∼ *N*^*ν*^, under various conditions. The solvent quality and polymer concentration-dependent *ν* (or 𝒟) are summarized in Table I.

**TABLE I:** Scaling exponent *ν* and fractal dimension 𝒟 of polymer chains under various conditions.

| Type | Solvent property | $\nu$ | $\mathcal{D}$ |
| --- | --- | --- | --- |
| Linear polymer in dilute solution | Poor | 1/3 | 3 |
| | Θ-condition | 1/2 | 2 |
|  | Good | 0.588 | 1.70 |
| Branched polymer in dilute polymer solution | Ideal | 1/4 | 4 |
| | Θ-condition | 7/16 | 2.29 |
|  | Good | 1/2 | 2 |
| Branched polymer in semidilute polymer solution | Semi-dilute (good solvent) | 0.395 | 2.53 |
| Branched polymer in dense polymer solution | Melts | 1/4 | 4 |

### Two length scales in neuron morphology

Individual neurons tend to feature densely branched structures, and hence their morphological characteristics can be analyzed in the language of polymer physics describing the conformation of branched polymers. Since branched polymers are composed of multiple sections of linear polymers, it is natural to consider two distinct ranges of length scale (*r*) (Fig. 1A) [32]: (i) The first range is defined at 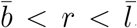. Here, 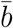 is the mean segment size defined between the data points in the reconstructed neurons, and 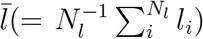 is the average length of linear sections (branches) between branch points or from one branch point to an axonal or dendritic tip. There is a great variation in length (*l*_*i*_) (Fig. 1B). The neurons over this scale are effectively described as a linear polymer. If the flexibility of a polymer in this length scale is significant, one could divide the polymer chain into multiple segments of Kuhn length; yet we have found that neurons in this length scale are typically stiff, lacking flexibility. (ii) The second range is defined at 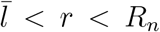, where *R*_*n*_ is the average size of the neuron, which is tantamount to the gyration radius 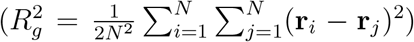 for a globular object (i.e., *R*_*n*_ = *R*_*g*_). In this range, the structural features of a neuron can be best represented by employing the structural characteristics of branched polymers.

**FIG. 1:**
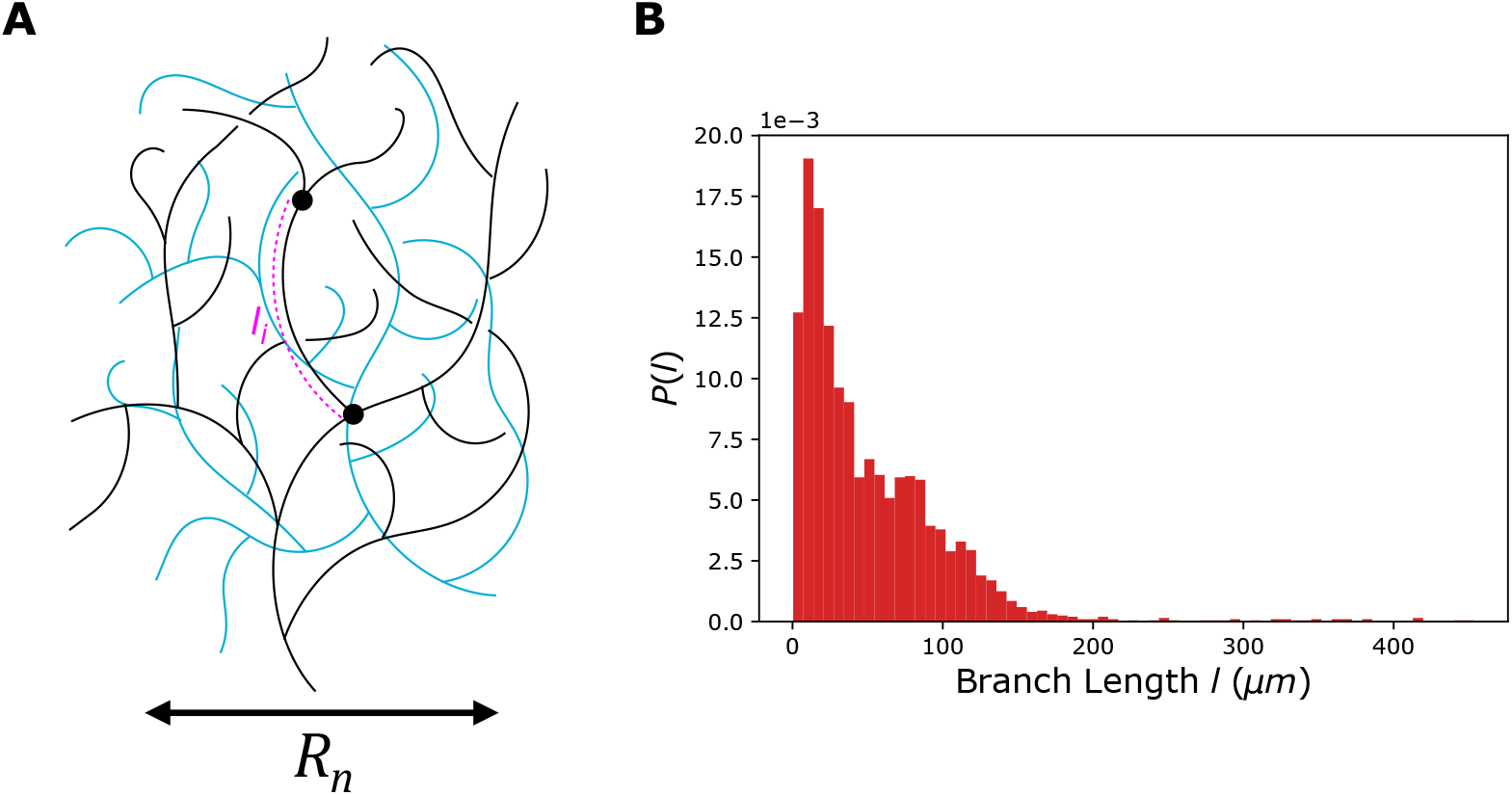
(A) Two different range of length scales associated with typical neuronal architecture mapped on a structure of branched polymer: (i) 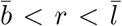 and (ii) 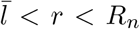. The black filled circles depict the branching points. *l*_*i*_ is the contour length of a branch, which is the segment between two branching points or between a branching point and a tip.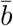 is the average size of segment defined between two consecutive data points along the neuron branch, 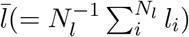 is the average length of branches, and *R*_*n*_ is the size of neuron. (B) The distribution of the branch length (*l*_*i*_) of neurons in the cluster 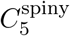 from the mouse V1.

### Form Factor

The form factor is routinely measured in scattering experiments to study the structure of materials [33–37]. Regardless of whether a given object is a polymer chain or a neuron, if the 3D coordinates of *N* monomers comprising the objects, {**r**_*i*_} (*i* = 1, 2, …, *N*) are available, one can numerically calculate the form factor using the following expression,

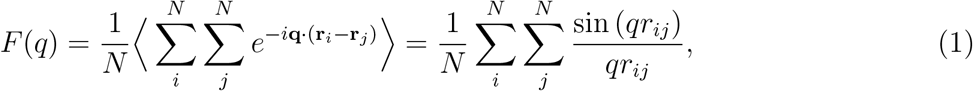

where **q** is the scattering wave vector, and **q** · (**r**_*i*_ − **r**_*j*_) = *qr*_*ij*_ cos *θ* with *q* = |**q**| and *r*_*ij*_ = |**r**_*i*_ − **r**_*j*_|. The second expression in Eq. 1 is obtained by averaging over the azimuthal angle *θ* between **q** and **r** − **r** and the polar angle *ϕ*, i.e., 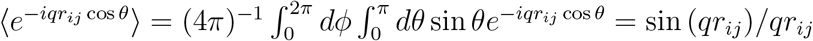. The wavenumber *q*, which can be considered as a resolution of the microscope, is inversely related to the length scale (*r*) to be probed (*q* = 2*π/r*), so that the *F* (*q*) at a high *q* regime probes an object locally, whereas *F* (*q*) at low *q* regime probes it globally.

Despite the numerical usefulness of the expression of *F* (*q*) given in Eq. 1, however, its physical meaning may not be immediately clear. In fact, in a physically more interpretable form, Eq. 1 is equivalent to the Fourier transform of distance distribution *p*(**r**) (see Method), namely

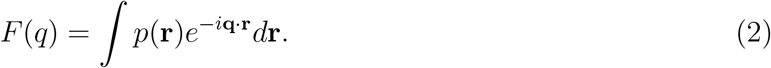

For 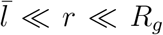 with *r* ≡ |**r**|, *p*(*r*) corresponds to the density of monomers inside a volume of a radius *r*, i.e., *p*(*r*) ≃ *n/r*^*d*^ where *n* is the average number of monomers inside the volume ∼ *r*^*d*^ at *d* dimensions. Provided that there is a scaling relationship between *n* and *r, r* ∼ *n*^*ν*^ with the scaling exponent *ν* (or the inverse fractal dimension 𝒟 = 1*/ν* that satisfies *n* ∼ *r*^𝒟^), which reflects how *n* monomers comprising the object span over the space defined by the length scale *r* [38], a dimensional analysis with *p*(*r*) ∼ *r*^1*/ν*−*d*^ and *d***r** ∼ *r*^*d*−1^*dr* in Eq. 2 yields [33]

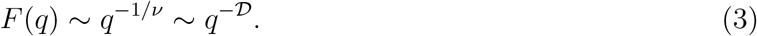

Depending on the wavenumber in the range of either 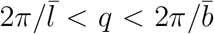 or 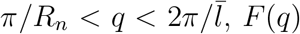, *F* (*q*) reveals different structures of neurons in terms of the scaling exponent *ν* (or 𝒟). On the other hand, for *qR*_*g*_ ∼ 1, corresponding to the Guinier regime, where the Fourier regime corresponding to the inverse of gyration radius is probed, *F* (*q*) is expressed as

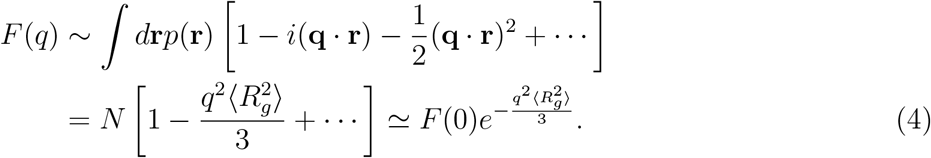

Thus, the radius of gyration of an object, 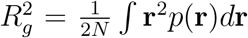, can be obtained from the slope of log *F* (*q*) vs *q*^2^ at small *q*, such that 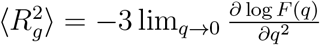.

Depending on the solvent quality and concentration of polymer solution as well as on whether the polymer chain is linear or branched, polymer chains adopt different conformations whose space-filling feature is captured in terms of the scaling exponent *ν*. In what follows, we review the scaling exponents of a linear or a branched polymer chain under various solvent conditions and at different concentrations of polymer solution [32, 33, 36].

### A polymer chain in dilute solution

According to the basic theory of polymer physics, the size (*R*_ℱ_, Flory radius) of a polymer chain scales with its length *N* as *R*_ℱ_ ∼ *N* ^*ν*^, where the scaling exponent *ν* changes with the solvent quality. In three dimensions (*d* = 3), a single flexible linear polymer adopts swollen conformations in a good solvent with *ν* = 0.588 but collapses to a globular form in a poor solvent with *ν* = 1*/*3 [33]. At the Θ-condition, which is a tri-critical point between a good and poor solvent condition where the attraction and repulsion at the level of two-body interaction compensate each other, a flexible polymer adopts conformation like those of ideal polymer that obeys the scaling law of *R*_ℱ_ ∼ *N* ^1*/*2^ [33, 39, 40]. Meanwhile, for a stiff polymer characterized by a large persistence length, it is expected that *R*_ℱ_ ∼ *N* with *ν* = 1 regardless of the solvent quality.

For 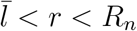, the neuron morphology can be studied by considering the structure of a branched polymer. To obtain the scaling exponent *ν* from *R*_ℱ_ vs *N* under various conditions, one can employ the Flory argument [32, 33, 36], which is often used to determine the solvent-quality dependent size of the polymer chain. In general, the Flory free energy of a polymer consisting of *N* monomers with the statistical segment of size *b* can be written as a function of polymer size *R*:

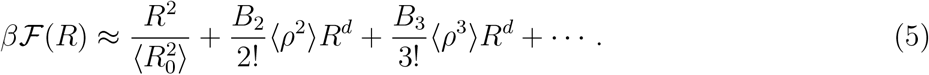

The first term is due to entropic contribution, 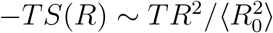, that arises from the elasticity of a polymer chain, which constrains the size of the polymer so that any attempt to deform the polymer engenders a restoring force of −*kR* with the spring constant of 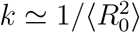, where 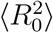 is the mean square size of an ideal polymer: 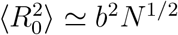 for randomly branched ideal polymers [36, 41], whereas 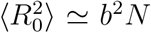 for linear polymers. The second, third, and remaining terms arise from the interaction between the monomers, *E*(*R*), whose effective density inside the pervaded volume of a polymer of size *R* is given as ⟨*ρ*⟩ ≃ *N/R*^*d*^. *B*_2_ is the second virial coefficient, which decides the strength of the two-body interaction; *B*_2_ > 0 is for the interaction between monomers in good solvent condition, making the two-body interaction repulsive, whereas *B*_2_ < 0 along with *B*_3_ > 0 delineates the poor solvent condition. Thus, the total free energy of a polymer chain results from the sum of entropic and energetic contribution ℱ(*R*) = −*TS*(*R*) + *E*(*R*), where the term −*TS*(*R*) tends to decrease *R*, whereas *E*(*R*) expands it. The size of the chain (Flory radius *R* = *R*_ℱ_) is determined by the balance between the two contributions, namely, by the minimization of Flory free energy, 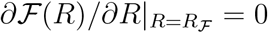.

With an assumption of zero correlation (⟨*ρ*^*n*^⟩ = ⟨*ρ*⟩^*n*^ = (*N/R*^*d*^)^*n*^), the Flory free energy is simplified as

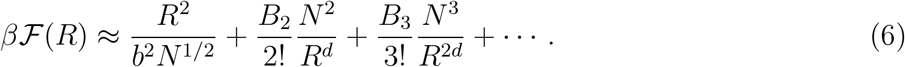

Under the good solvent condition, the two-body interaction is more dominant than the higher-order terms, and the free energy can be truncated up to the two-body interaction term, allowing us to determine the *R*_ℱ_ in a good solvent,

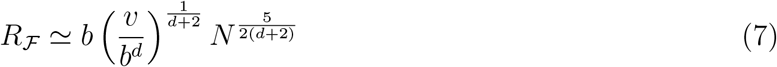

with *B*_2_ = *v*. Thus, the size scaling exponent of a randomly branched polymer in a good solvent at *d* = 3 is 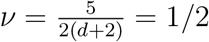 (i.e., 𝒟 = 2) [32].

Under the Θ-solvent condition, the second virial coefficient vanishes (*B*_2_ = 0), and the Flory radius of a branched polymer is determined from the balance between the elastic free energy and the three-body interaction term with *B*_3_ = *w* in Eq. 5 as

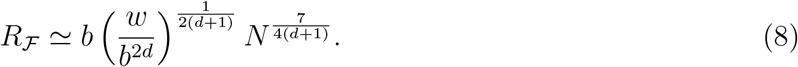

Thus, the size scaling exponent of a randomly branched polymer under Θ-solvent condition at *d* = 3 is *ν* = 7*/*16 (𝒟 = 1*/ν* = 2.29) [32].

### A polymer chain in semi-dilute/dense polymer solution

When the concentration of polymer solution increases, the individual chains start overlapping, influencing the chain conformation of neighboring polymers, which defines the overlap volume fraction Φ_*c*_ ∼ *Nb*^3^*/R*_ℱ_ ^3^ ∼ *N* ^1−3*ν*^. In the semi-dilute regime (Φ_*c*_ ≲ Φ ≪ 1), it is no longer discernible whether two spatially adjacent monomers are from the same chain or different chains, and the original strength of the intra-polymer two-body interaction (*B*_2_ = *v*) is screened to yield *B*_2_ ≃ *v/N* ^1*/*2^. In this case, the Flory free energy is written as

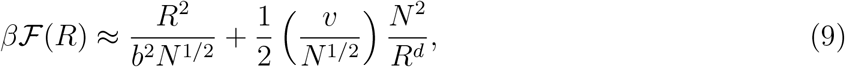

and 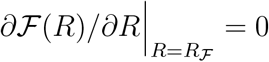 yields the Flory radius of

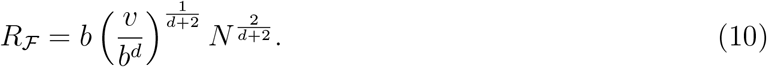

Thus, the size scaling exponent of a randomly branched polymer in a semi-dilute polymer solution under good solvent conditions is *ν* = 2*/*(*d* + 2) = 2*/*5 = 0.4. This number is close to the exponent *ν* = 0.395, the inverse fractal dimension of percolating clusters at percolation threshold (*p* = *p*_*c*_), 𝒟 = 1*/ν* = 2.53, estimated from computer simulations [42].

For dense melts of branched polymers (Φ_*c*_ ≪ Φ ≈ 1), the interactions between polymer segments are fully screened. In this case, individual branched polymers behave as if they are effectively in an *ideal* condition, giving rise to the scaling relationship of *R*_ℱ_ ∼ *N* ^1*/*4^, and interesting scaling *F* (*q*) ∼ *q*^−4^ emerges in the range of intermediate 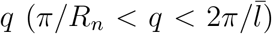. Since the linear sections of neurons 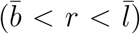 are stiff and effectively rigid rod-like, the length of the statistical segment (Kuhn length) is 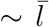, and the number of statistical segments comprising the branched polymer is 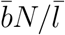. Therefore, the size of polymer scales as 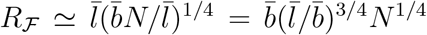. The fractal dimension of 𝒟 = 4 may sound geometrically impossible; however, an object with fractal dimension 𝒟 = 4 is still permissible in the range of 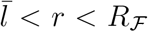 because the stiff linear section creates extra space at 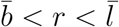 [36].

Note that a scaling behavior of *F* (*q*) ∼ *q*^−4^ could also emerge at a large *q* regime (*q* ≳ *b*^−1^), although it is not directly relevant to our problem that discusses branched polymer-like neurons comprised of rigid segments with large persistence length. This corresponds to the Porod scattering 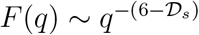 with 𝒟_*s*_ = 2 [43–45], contributed by smooth surfaces of compact globules (𝒟_*s*_ = 2) that are formed by a *flexible* chain under a poor solvent condition.

Although neurons are without looped topology, the Flory argument, demonstrated in the foregoing two subsections, can be used to derive the scaling relations of polymer chains with closed-loop structures as well. The looped chain objects are smaller in size, affecting the pre-factor of scaling relation; yet their scaling exponents remain unaltered from those of linear objects as long as the solvent quality is unchanged.

The scaling exponents (fractal dimensions) for different types of polymers are summarized in Table I.

## RESULTS

Here we present our *F* (*q*)-based analysis on the datasets of neuron morphology reconstructions from three distinct organisms with an increasing degree of complexity (Fig. 2): (i) the *C. elegans* nervous system, (ii) projection neurons in the *Drosophila* olfactory system, and (iii) mouse primary visual cortex neurons.

**FIG. 2:**
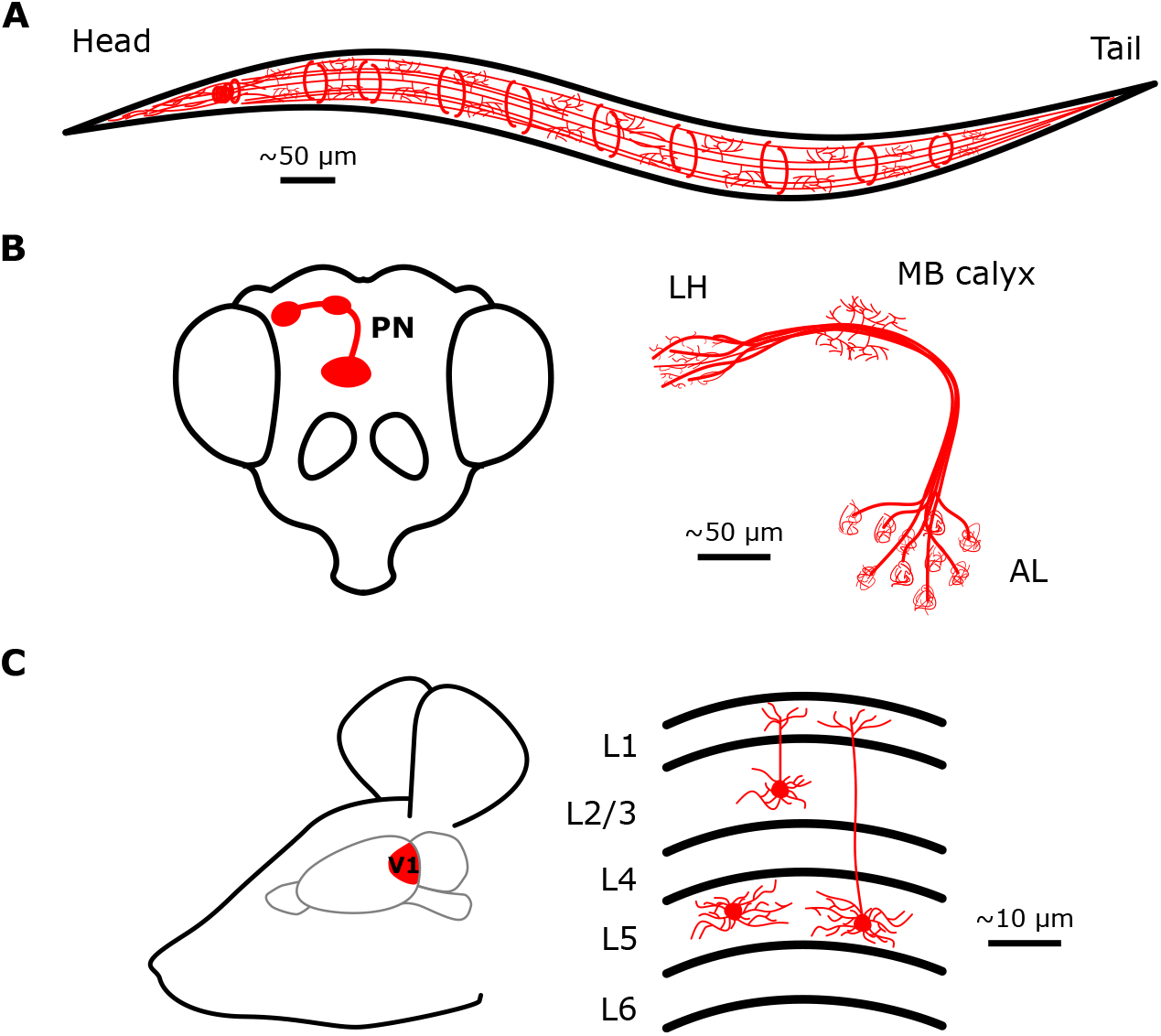
Diagrams of three systems where *F* (*q*)-based analysis was made: (A) the *C. elegans* nervous system, (B) *Drosophila* olfactory projection neurons, and (C) Mouse primary visual cortex.

### Neurons in *C. elegans*

*C. elegans* is one of the organisms whose neural connectivity has been fully mapped. Morphological reconstruction of the *C. elegans* nervous system is available as a part of the OpenWorm initiative which created a complete connectome of the organism [31] (see Fig. 2A). *C. elegans* has 302 neurons with their morphologies and inter-neural connectivity being fully specified [46, 47]. Because of its relative simplicity, it is an ideal organism to explore the basis of neural dynamics and brain function [48–50]. Partial, but more detailed reconstructions are also available for small parts of the sensory neurons [51–53]. Complex neural behaviors of both associative and non-associative learning are realized through these neurons [47] which are functionally classified into sensory, motor, interneurons, polymodal, and unknowns [31, 47, 54, 55] (Fig. 3). Respective circuits for specific behaviors such as locomotion and chemosensory responses are well-documented as well [47, 49, 52, 56].

**FIG. 3:**
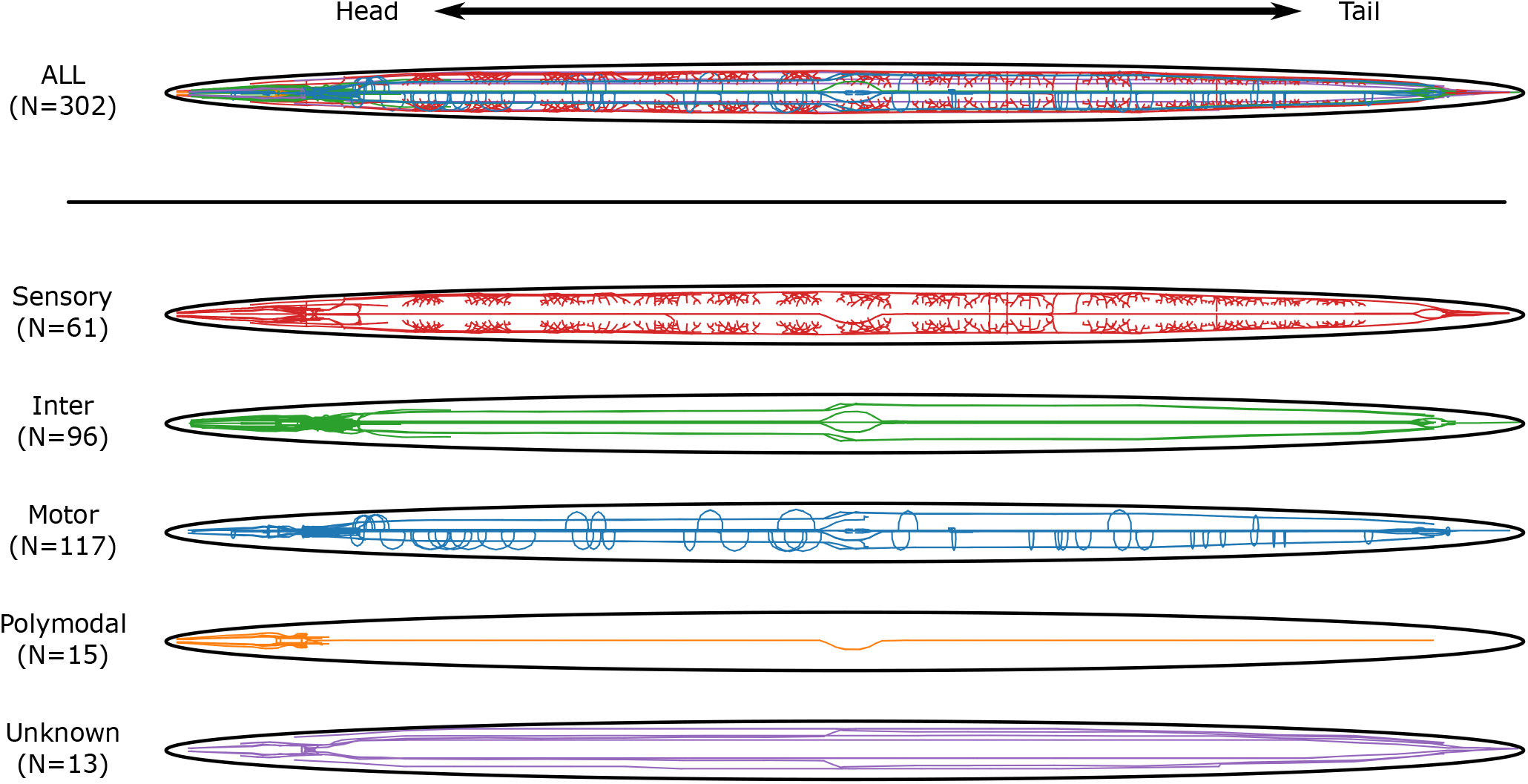
Illustrations depicting the morphological reconstructions of neurons in *C. elegans*. Depicted at the top is the entire *C. elegans* nervous system. The nervous systems are redrawn for the neurons of different functional types.

The overall architecture of neurons in the *C. elegans* nervous system largely conforms to the organism’s tubular body shape, which is reflected in the *F* (*q*) plots of large neurons classified into clusters 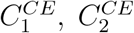, and 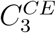. The neurons in these clusters exhibit 𝒟 ≈ 1 at the wavenumber smaller than the dips formed at *q* ≈ 2*π/d*_*C*_(≈ 0.13 *μ*m^−1^) (dotted lines in Fig. 4A) where *d*_*C*_(≈ 47.9 *μ*m) is the average diameter of the cross-section [57, 58]. The 2/3 of neurons (*N* = 198) come in a pair projecting on the left and right sides of the body, and the rest (*N* = 104) are largely confined either in the head or in the tail region.

**FIG. 4:**
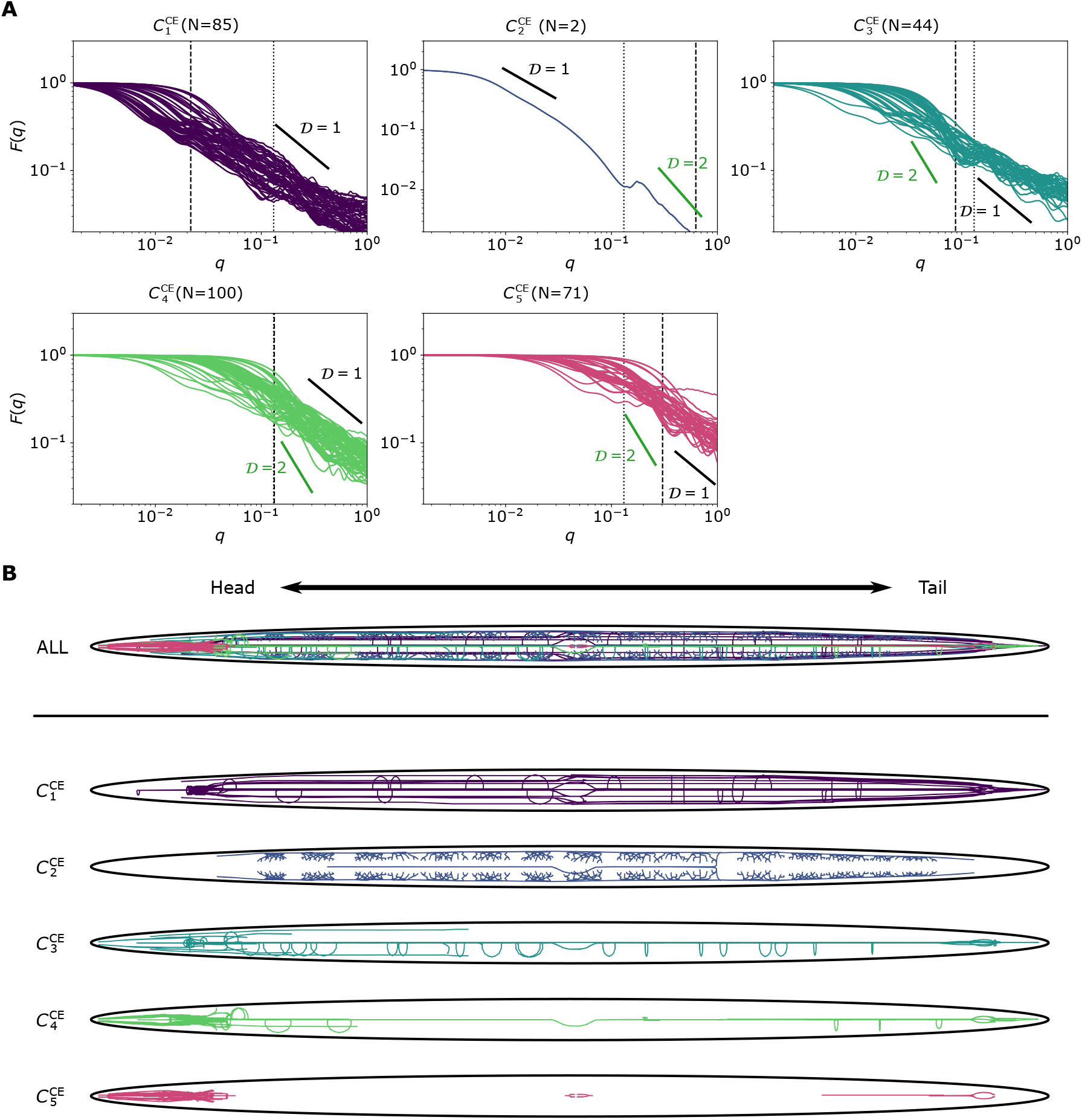
(A) *F* (*q*) plots of neurons in the *C. elegans* nervous system grouped by the *F* (*q*)-based clustering result. The dotted lines denote *q*(= 2*π/d*_*C*_) corresponding to the inverse scale of the average diameter (*d*_*C*_) of adult *C. elegans*. The dashed lines denote 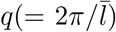 correspond to the average branch size 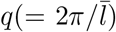 of neurons in the cluster. Reference fractal dimensions (𝒟s) are highlighted in the *F* (*q*)-plots with the slopes. The clusters from 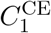 to 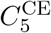 are indexed based on the average size (*R*_*n*_) of neurons in each cluster in a decreasing order. (B) Illustrations depicting the morphological reconstructions of neurons in *C. elegans* grouped by the *F* (*q*)-based clustering. The same colors are used as in (A) for each cluster.

The *F* (*q*)-based clustering groups 302 neurons into five clusters based on their overall morphological features (see Fig. 4 and Fig. 5). (i) The neurons involved with the interneurons (*N* = 34) and motor function (*N* = 37), which span across the entire body, are grouped into the cluster 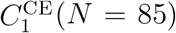. The rest of the cluster is composed of 6 sensory neurons, a polymodal neuron, and 6 unknowns. (ii) The cluster 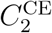 is made of 2 PVD neurons (PVDL and PVDR), which are known to be responsible for the sensory detection of drastic changes in mechanical forces, osmolarity, and temperature [59–61]. They feature a dendritic morphology of exceptional branching patterns repeating across the entire body (Fig. 4B). The *F* (*q*) of the PVD neurons scales *F* (*q*) ∼ *q*^−2^, exhibiting the fractal dimension of 𝒟 = 2 at 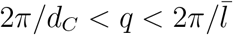 (Fig. 4A), indicating that the morphology of PVDs with intricate branching pattern (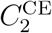 of Fig. 4B) is similar to the configuration of a self-avoiding branched polymer in dilute solution. (iii) The neurons in cluster 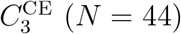, structurally similar to those in 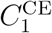 but are shorter in branch length 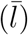, are mostly motor neurons (*N* = 32). The cluster also contains a small number of sensory (*N* = 5) and interneurons (*N* = 7). (iv) The clusters 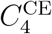 and 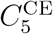 are comprised of small to mid-sized neurons mainly confined around the head region. Unlike the other neuron clusters, the neurons grouped in cluster 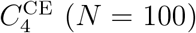 are diverse in terms of neuron type composition, containing 44 sensory neurons, 22 interneurons, 22 motor neurons, 6 polymodal neurons, and 6 unknowns. The cluster 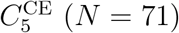 consists of 4 sensory neurons, 33 interneurons, 26 motor neurons, and 4 polymodal neurons. The clusters 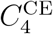 and 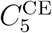 can be further classified into smaller clusters by repeating the application of the same clustering algorithm to each cluster (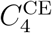 and 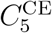 in Fig. 5A, and Figs. 5B and 5C).

**FIG. 5:**
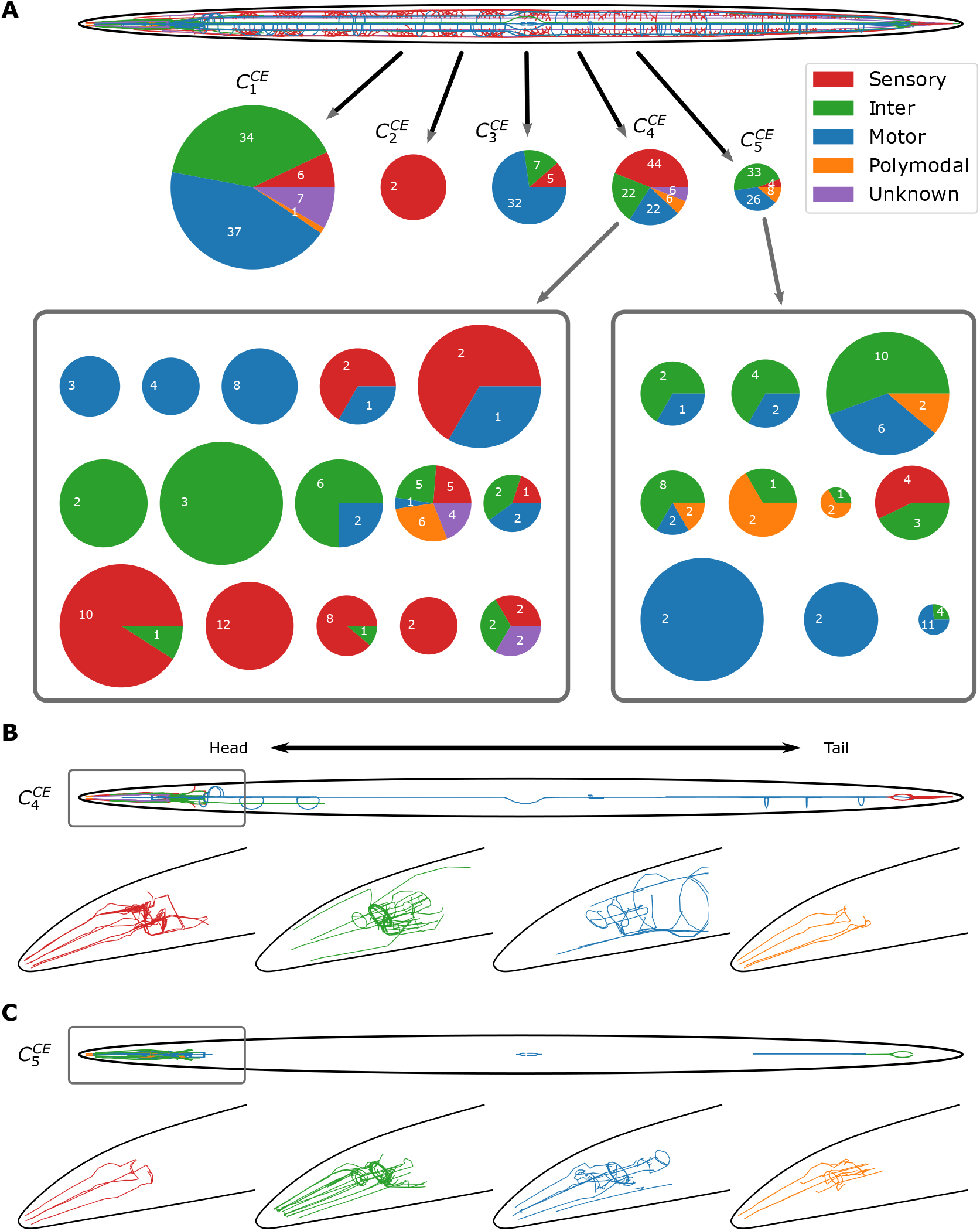
(A) Pie charts depicting the functional composition of *F* (*q*)-based clusters in the *C. elegans* nervous system. The subclusters of 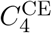 and 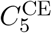 when the same clustering algorithm is applied repeatedly are shown inside the boxes. The size of the pie charts reflects the total contour length of neurons in each cluster. Illustrations depicting the morphological reconstructions of neurons in (B) the cluster 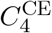 and (C) the cluster 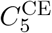 with functional labeling. The enlarged 3D reconstruction of neurons in the head region, marked by gray boxes, is depicted below for each functional type.

Taken together, it is found that the motor and interneurons grouped in 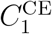 and 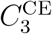 extend across the entire body of *C. elegans*, lacking branches. In contrast, the neurons with sensory function (PVDs) grouped in 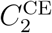 are characterized by the pattern of extensive branching. Although the neurons classified in the clusters 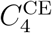 and 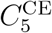 are still in a mixture of many neurons with different functional types, their sizes (contour lengths) are smaller than those in 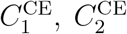, and 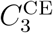. Overall, the *F* (*q*)-based clustering, which captures the morphological features of neurons, has grouped the neurons in the *C. elegans* nervous system into their respective functional type to a first approximation.

### Projection neurons in three neuropils of *Drosophila* olfactory system

The full *Drosophila* hemibrain connectome constructed from electron microscope images [13, 62] offers neuron morphologies at high resolution. The neurons constituting the second-order layer of the *Drosophila* olfactory system are made of three types of projection neurons, which include uniglomerular PNs (uPNs), multiglomerular PNs (mPNs), and local neurons (LNs). Among them, uPNs (*N* = 111) are bundled together into on average 3 uPNs and comprise ∼ 50 distinct glomeruli, which receive signals from ∼ 50 different types of olfactory receptor neurons (ORNs) where a diverse array of chemical signals are encoded via the combinatorial coding [63–67]. The uPNs extend across the layer of second-order neurons, bridging between ORNs and higher olfactory centers (see Fig. 2B). The multiglomerular PNs (mPN) (*N* = 30) extend over multiple glomeruli, and the local neurons (LNs) (*N* = 12) are confined within the neuropil.

The PNs extend their dendritic and axonal branches and these branches are densely entangled to form three neuropils. Along the neural signal transmission pathway, the antennal lobe (AL) is the first neuropil, consisting of the ∼ 50 glomeruli, and the axonal extensions of the uPNs in AL project to the other two neuropils, mushroom body (MB) calyx and lateral horn (LH), which are the sites that synapse with Kenyon cells and lateral horn neurons for learned and innate responses to olfactory signals, respectively [68–72]. Since neuron projections in each neuropil feature unique morphology which is discernible from one neuropil to another, we segmented the uPNs into the parts contributing to each of the three neuropils [73] and analyzed them separately.

In AL, *F* (*q*)-based clustering results in seven clusters (Fig. 6), characterized by 2 ≤ 𝒟 ≤ 4 at 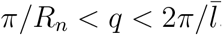. The clusters 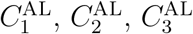, and 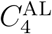, which represent the major population of the 127 PN projections to AL, are characterized by the structure of a branched polymer in melts (𝒟 = 4), semi-dilute polymer solution (𝒟 = 2.53), and dilute polymer solution under Θ-solvent (𝒟 = 2.29) or good solvent (𝒟 = 2) condition.

**FIG. 6:**
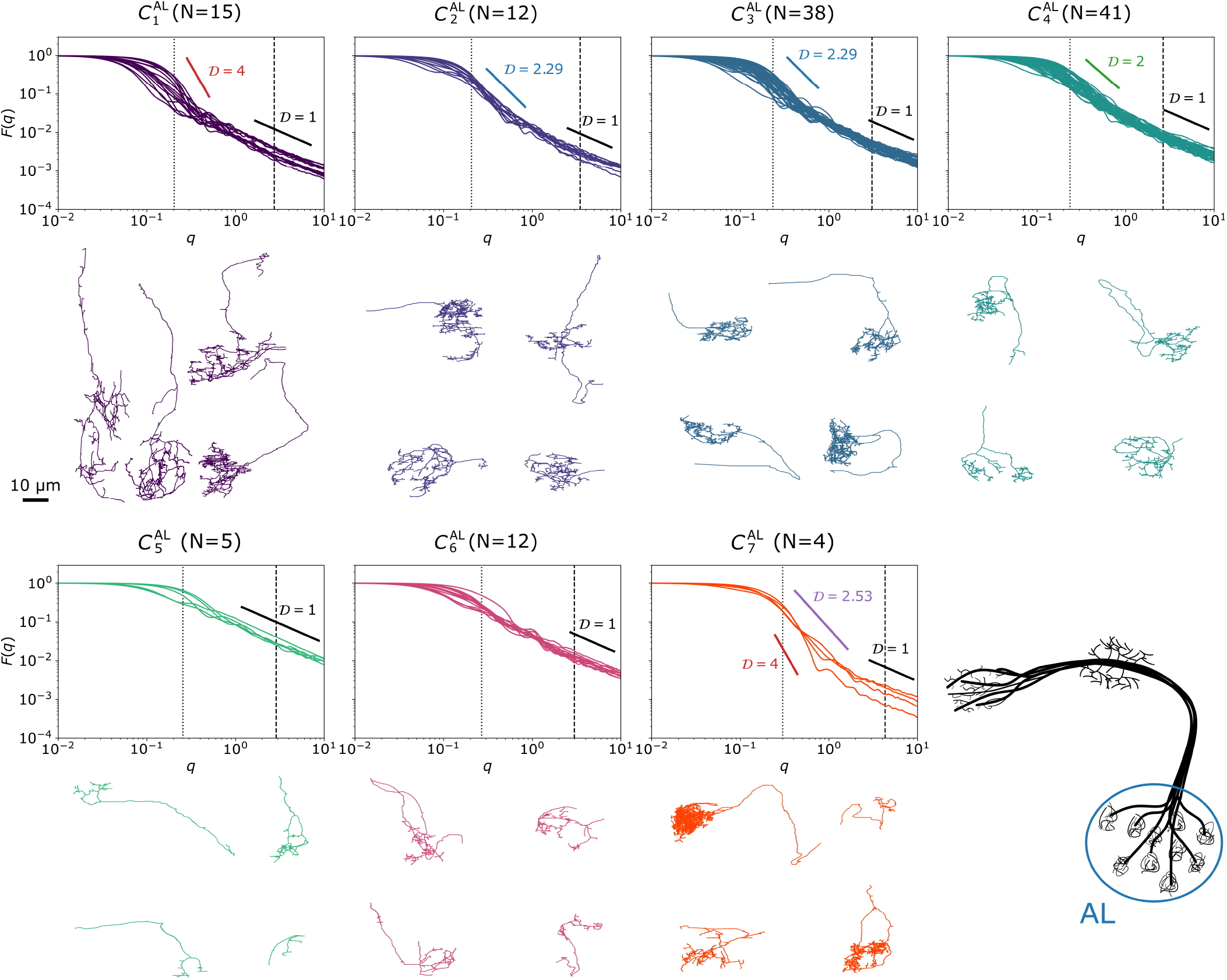
*F* (*q*) plots of the PN projections to AL in the *Drosophila* olfactory system grouped by the *F* (*q*)-based clustering result. The dotted lines denote *q* corresponding to the inverse scale of neuron size (*q* = *π/R*_*n*_) of the cluster, and the dashed lines correspond to the average branch size 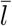 of neurons in the cluster 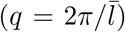. Reference fractal dimensions (𝒟s) are highlighted in the *F* (*q*)-plots with the slopes. The clusters from 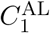 to 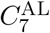 are indexed based on the average size (*R*_*n*_) of neurons in each cluster in a decreasing order.

In MB calyx, *F* (*q*)-based clustering identifies six clusters (Fig. 7). The minor cluster *C*_1_ is comprised of 3 regulatory neurons. These neurons are either an MB output neuron (MBON) or an MB-C1 (inhibitory interneurons) [13], whose shapes greatly differ from typical PNs. Remarkably, the *F* (*q*)-based clustering correctly separates them. In the clusters *C*_2_, *C*_3_, *C*_4_, and *C*_5_, the PNs exhibit boutons of the size around 3 − 6 *μ*m [74] at the tip of protrusions stemming from the main axon (Fig. 7). KCs form claw-like projections and envelope boutons at the synaptic site [75]. The structural characteristic of the two-layered hierarchy manifests itself in the *F* (*q*) plots with 𝒟 ≈ 2.29, indicating that axonal branches adopt the configurations of branched polymer in the Θ-solvent at two different length scales, below and above the ‘plateau’ at *q* ≈ 0.4 *μ*m^−1^ (*r* ≈ 16 *μ*m) (see the arrows pointing the plateau of *C*_2_ and *C*_3_ in Fig. 7). This trend is practically absent in clusters *C*_5_ and *C*_6_ which have only a small number of boutons in each PN.

**FIG. 7:**
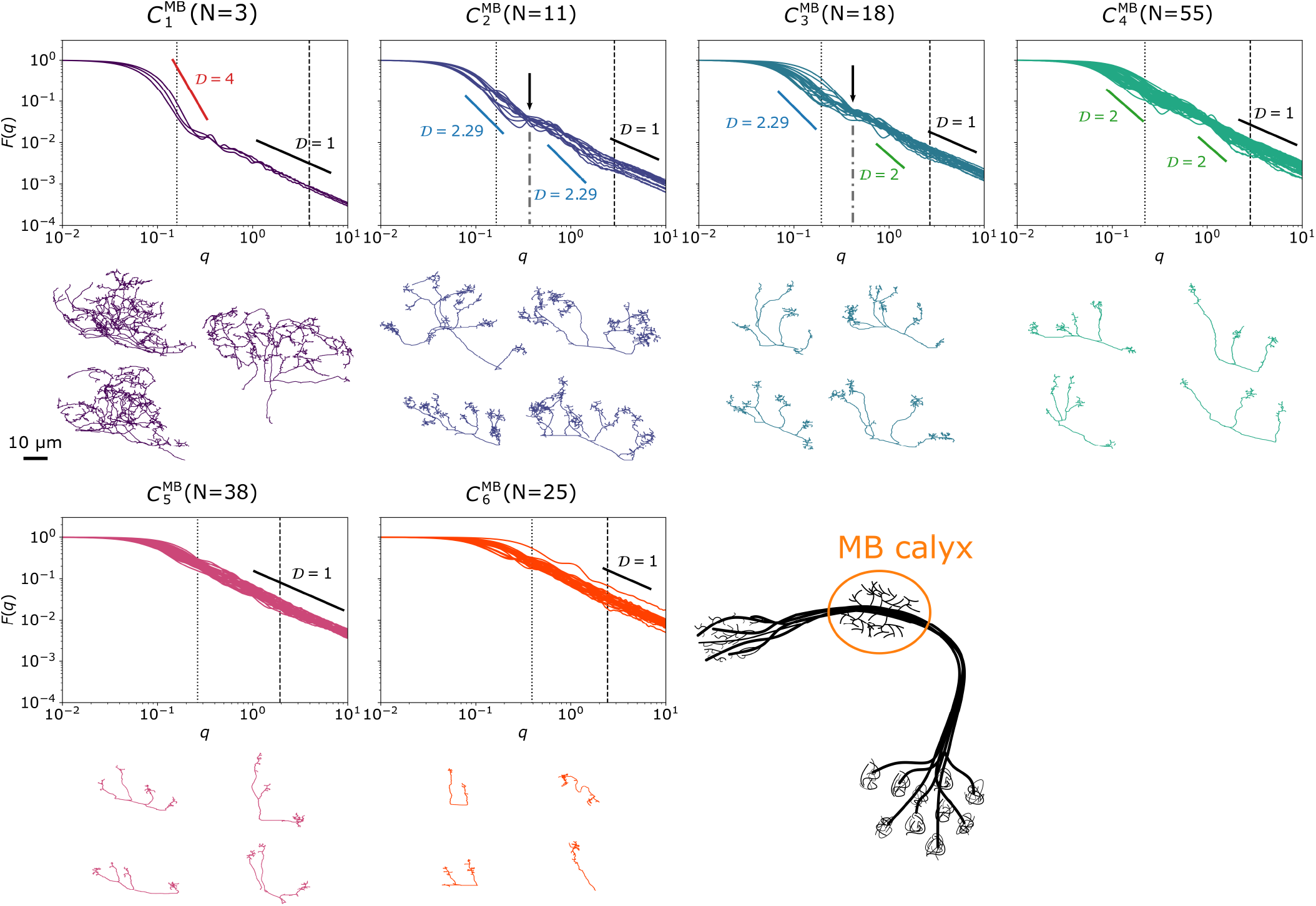
*F* (*q*) plots of the PN projections to MB in the *Drosophila* olfactory system grouped by the *F* (*q*)based clustering result. The locations of plateau in *C*_2_ and *C*_3_ at *q* ≈ 0.4 *μ*m^−1^ are indicated by the arrows. Other details are the same as Fig. 6.

In LH, there are six clusters (Fig. 8). Similar to *C*_2_ and *C*_3_ in MB calyx, the clusters 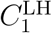 and 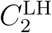 in LH are characterized by the *F* (*q*) displaying a ‘plateau’. Despite the overall similarity in the shape of *F* (*q*) in that a plateau is present at intermediate *q*, the details of *F* (*q*) and the actual PN morphologies in LH are slightly different from those in MB calyx (Fig. 8). The plateaux of 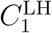 and 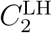 are identified at *q* ≃ 0.5 − 0.6 *μ*m^−1^ (*r* ≃ 11 *μ*m). Furthermore, visual inspection of the morphologies indicates that smaller axonal branches are uniformly distributed over the main branches of PNs in LH. From 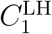 to 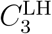, the PNs change their morphologies from 𝒟 = 4 to 𝒟 = 2.29 and 𝒟 = 2. In the remaining clusters, 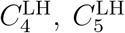, and 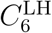, PNs are small in size and relatively featureless (𝒟 = 1). The difference between the clusters stems from the overall size (*R*_*n*_).

**FIG. 8:**
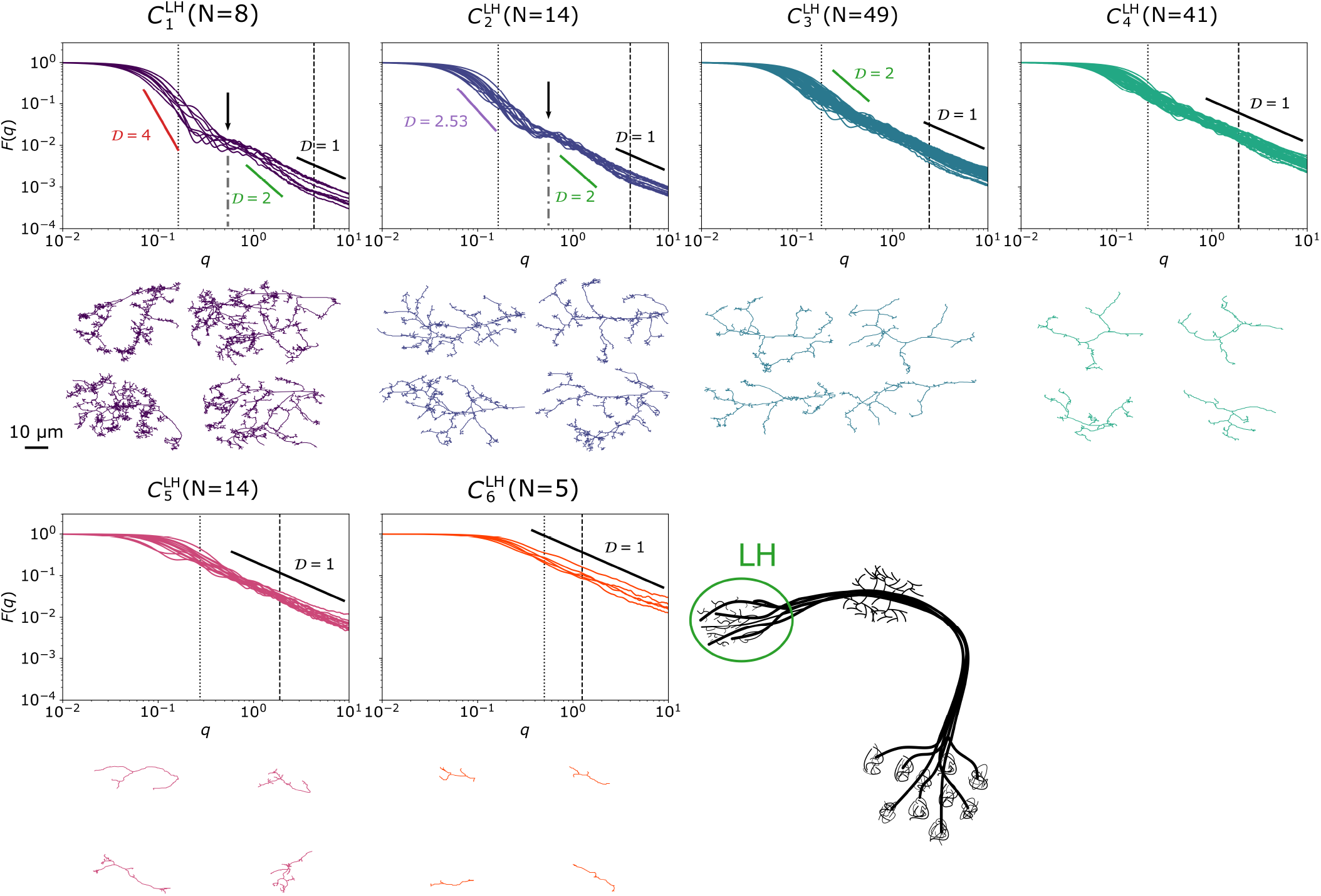
*F* (*q*) plots of the PN projections to LH in the *Drosophila* olfactory system grouped by the *F* (*q*)based clustering result. The locations of plateau in 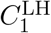 and 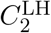 at *q* ≈ 0.5 − 0.6 *μ*m^−1^ are indicated by the arrows. Other details are the same as Fig. 6.

In the second-order neurons of the *Drosophila* olfactory system, PNs with different morphology are mixed to constitute synaptically dense MB calyx and LH.

### Neurons in mouse primary visual cortex

The Allen Cell Type database contains full and partial reconstructions of neurons in the mouse primary visual cortex (V1), which plays a critical role in visual processing as the first cortical region to receive visual input [76]. Visual signals processed in V1 are sorted [77] and relayed to at least 18 different brain areas [78–80] which are deemed functionally distinct [81]. The V1 is divided into 6 functionally distinct layers [82]. For example, layers 2 and 3 house many direction-selective oriented cells [83]. Layer 5 is known for mostly non-oriented cells with a large receptive field [83]. Diverse types of neurons with varying sizes spanning across multiple layers (see Fig. 2A) comprising V1 are rich in pyramidal cells, whose organization maps the orientation preferences [84]. Gouwens *et al*. performed hierarchical clustering on the Allen Cell Type database using a combination of morphometric features such as branch number and the total contour length of the neurons [11]. All neurons in the Allen dataset are categorized under two different dendrite types, either spiny or aspiny based on the existence of dendritic spines, which are small protrusions along the dendrite where synaptic inputs occur. Although individual neurons are labeled with dendrite type information, these minuscule structures are not represented in the morphological reconstructions. Here, instead of relying on morphometry, we calculated the *F* (*q*)s of V1 neurons by leveraging the 3D coordinates provided in the database to assess the similarity between them and repeated Gouwens *et al* ‘s clustering procedure.

In comparison with Gouwens *et al*. who classified V1 neurons into 19 clusters for spiny neurons and 19 clusters for aspiny neurons [11], our *F* (*q*)-based clustering produced 11 and 8 clusters for spiny and aspiny neurons, respectively (see Fig. 9, Fig. 10, and visit https://github.com/kirichoi/ℱqClustering for the complete neuron morphologies per cluster). The difference in the number of clusters likely stems from the fact that information on relative soma depth, for instance, utilized in morphometry-based clustering, is not explicitly included in the *F* (*q*)-based clustering.

**FIG. 9:**
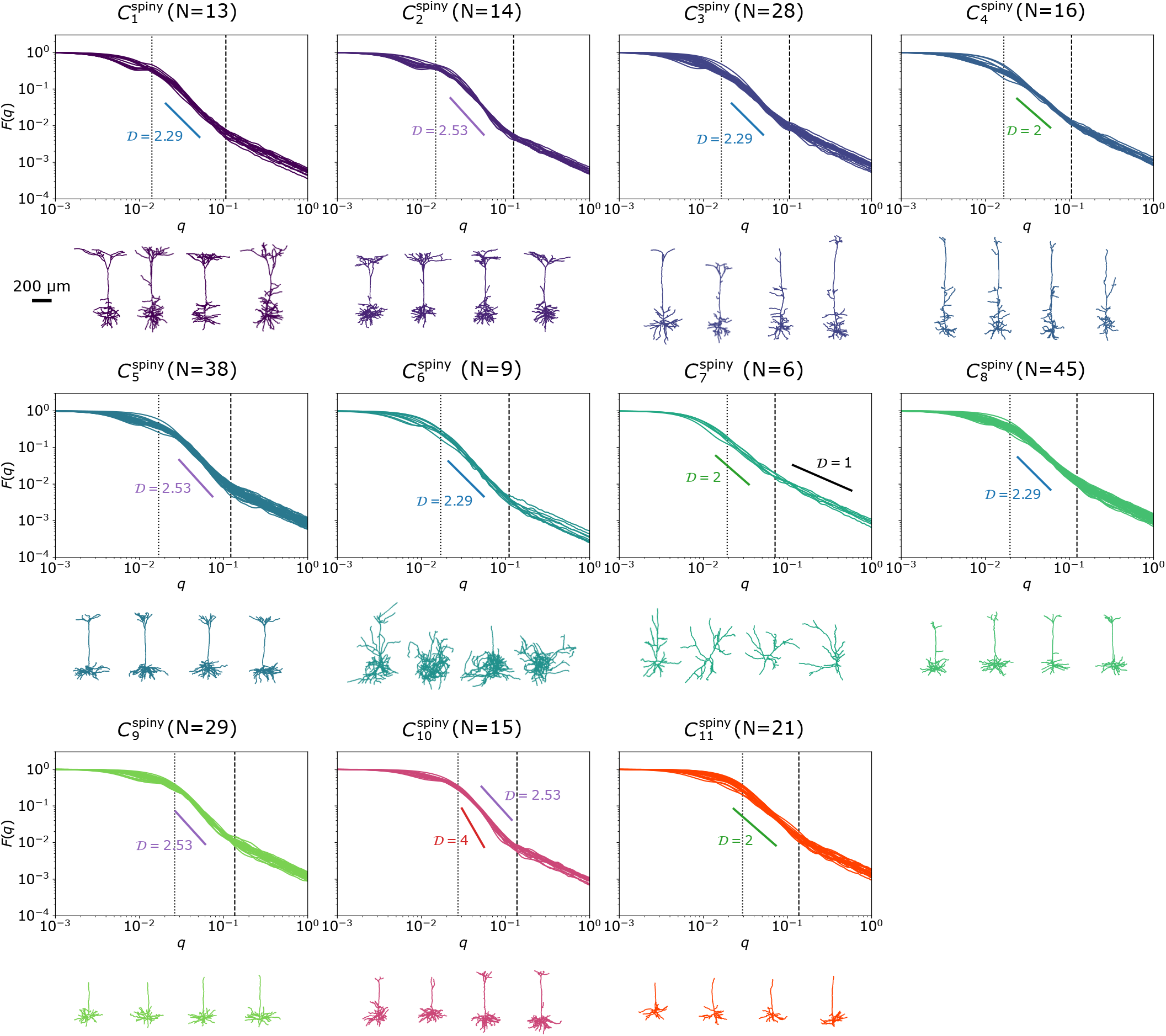
*F* (*q*) plots of spiny neurons in the mouse primary visual cortex grouped by the *F* (*q*)-based clustering result. Other details are the same as Fig. 6.

**FIG. 10:**
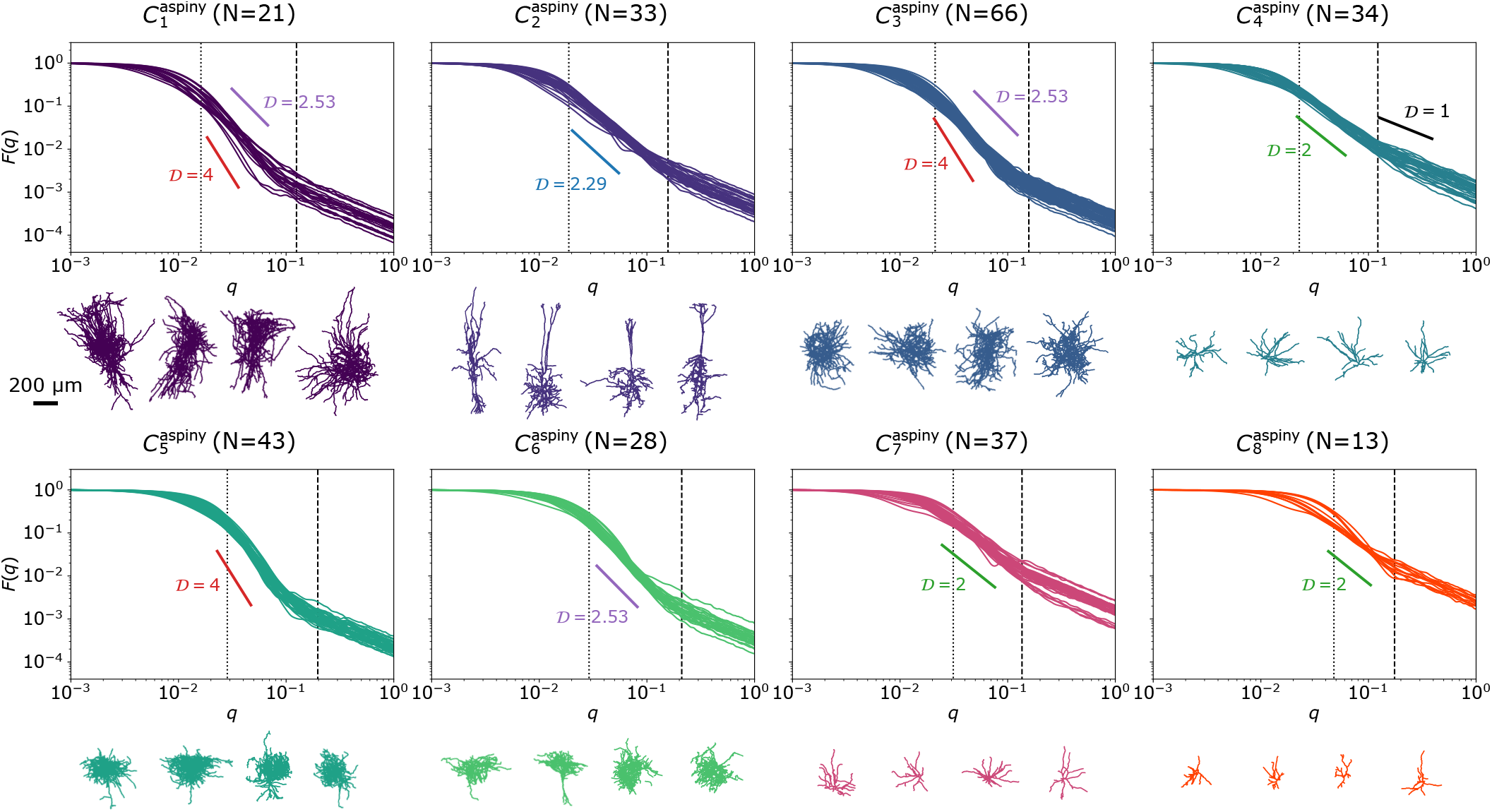
*F* (*q*) plots of aspiny neurons in the mouse primary visual cortex grouped by the *F* (*q*)-based clustering result. Other details are the same as Fig. 6.

From the *F* (*q*) plots, it becomes clear that neurons are classified based on 𝒟 and *R*_*n*_ into several distinct subgroups. For spiny neurons, clusters 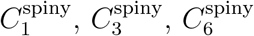, and 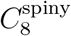 all share 𝒟 = 2.29 for 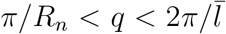 (or 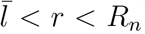) (Fig. 9). The clusters 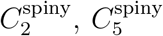, and 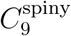 are featured with 𝒟 ≈ 2.53. Morphologically, these clusters exhibit relatively denser dendrites with more number (or higher density) of branches than those clusters with 𝒟 = 2.29 (Fig. 9). The clusters 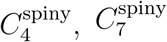, and 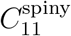 display the same fractal dimension, 𝒟 = 2, at 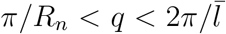; yet differ from each other by their neuron size 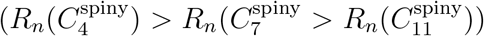. The cluster 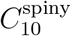 displays 2.53 < 𝒟 < 4, featuring unusually dense arborization, similar to that of a branched polymer in a melt. For aspiny neurons (Fig. 10), we find that the clusters 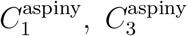, and 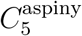 are characterized with 𝒟 = 4; 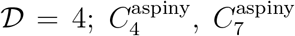, and 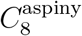 with 𝒟 = 2; 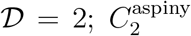 with 𝒟 = 2.29; and 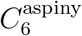 with 𝒟 = 2.53.

We notice that the spiny neurons are on average characterized with 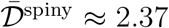, whereas the aspiny neurons with 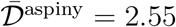. Thus, the neuron mass density (*ρ* ∼ *n/r*^3^ ∼ *r*^𝒟−3^ ∼ *n*^1−3*/*𝒟^) is higher in aspiny neurons 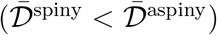. It is noteworthy that the absence (or presence) of apical dendrites, whose size is comparable to the size of a neuron (*R*_*n*_), does not contribute significantly to the fractal dimension in the intermediate scale 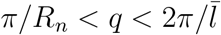.

Compared with the morphology types defined by Gouwens *et al*., *F* (*q*)-based clustering prioritizes the overall structural similarity dictated by 𝒟 and *R*_*n*_ (and more precisely by *d*_*ij*_ of Eq. 12). Still, the similarity between the two clustering outputs is statistically meaningful, as the Pearson’s *χ*^2^test of independence between the two clustering outputs results in very small p-values for both the spiny and the aspiny neurons (*p* = 9.368 × 10^−33^ and *p* = 8.736 × 10^−21^ respectively). Other similarity metrics that enable us to compare the two clustering outputs, such as the Baker’s Gamma index [85] 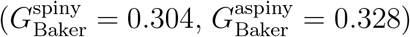, normalized mutual information [86] (NMI_spiny_ = 0.355, NMI_aspiny_ = 0.302), and *v*-measure [87] (*v*_spiny_ = 0.355, *v*_aspiny_ = 0.302) all point to a moderate similarity between the two clustering outputs. The *F* (*q*)-based clustering correctly captures the morphological characteristics of the neurons and can be utilized as an alternative criterion for clustering.

## DISCUSSION

The *F* (*q*)-based clustering produces a comprehensible and systematic classification of neurons, complemented by the size (*R*_*n*_) and the fractal dimension (𝒟) that quantifies the extent of the branching pattern of reconstructed neurons. We argue that different groups of neurons classified based on their morphology has functional significance. In particular, the branchness of neurons, quantified by 𝒟, is of great significance to the firing activities of neurons, which should be determined by the densities of ion channels on the membrane surface. For a neuron characterized with 𝒟, it is expected that the conductance (*G*) per neuron is proportional to the total number of ion channels (*n*_ch_) or the capacity of the neuron dictated by the total contour length 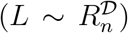. If one imagines that a neuron is locally a cylindrical object with a cross-sectional diameter *d*(*x*) at a given position *x* along its contour [28] and that the density (*σ*_ch_) of ion channels on the membrane surface is given, the conductance of ions across the membrane surface of a neuron is estimated as

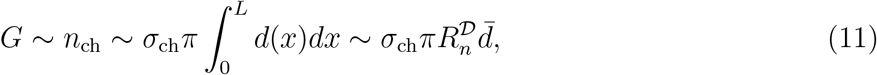

where 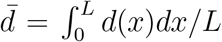 is the average cross-sectional diameter of a neuron. A densely branched neuron characterized with a greater fractal dimension, say 𝒟 = 4, is expected to display a higher conductance than a branchless neuron with 𝒟 = 1. Functional differentiation of neurons, dictated by various firing patterns, has indeed been associated with a morphologically heterogeneous neuronal population [3].

The clustering based on topological characteristics (e.g. morphometrics) [11, 15, 17–19] is conceptually akin to our *F* (*q*)-based clustering. Although the morphometrics-based classification could be more comprehensive with a set of geometrical features selected, it requires a more extensive preprocessing as well as careful selection of feature sets to avoid introducing a bias or redundancy. It is not our intention to argue that our *F* (*q*)-based analysis is superior to other metric-based analyses, but it can still be said that our *F* (*q*)-based analysis is more objective in that it relies on the 3D coordinates of neurons with no prior knowledge of the neuron structure. Accompanied by electrophysiological measurement [10, 11] and stimuli response profiles [88], our *F* (*q*)-based classification holds good promise for a systematic study to elucidate the structure-function relationship in neurons, namely, the relationship between the neuron morphologies and the neural activities.

## METHODS

### Data preparation

- A total of 302 morphological reconstructions of neurons in the *C. elegans* nervous system were collected from the OpenWorm dataset [31]. The neuron type information available from the dataset was used to assign each neuron into one of the five functional categories (sensory, motor, interneuron, polymodal neuron, and others).
- For the *Drosophila* olfactory projection neurons, we use the TEMCA2 dataset [13]. A total of 162 *Drosophila* olfactory neurons are present in the dataset, which is extracted from the right hemisphere of the female *Drosophila*. From the dataset, three neurons (ID=203840, 2738003, 2738059) that are identified to be not PNs and six neurons (ID=1549518, 2738042, 2738083, 2738261, 2970058, 2970073) that did not project to any of the neuropils are dropped. We removed artifacts in the data in which certain data points are dramatically offset from the expected trajectories due to misalignments of images when constructing the original morphological skeleton. These data points were identified by a geometrical criterion that the distances between three consecutive points should be smaller than 10 *μ*m. Next, we systematically segmented the neurons inside the three neuropils using the same process detailed in [73].
- The Allen Cell Type Database [11] contains 509 full and partial morphological reconstructions of neurons in the mouse primary visual cortex. Following Gouwens *et al*. [11], we classify the whole neurons into 234 spiny and 275 aspiny neurons. When comparing the clustering results, we noticed that Gouwens *et al*. used 461 morphological reconstructions which is smaller than what is currently available in the database. Gouwens *et al*. provided the exact cell IDs for 461 morphological reconstructions, but a subset of cell IDs do not match with those currently present in the database. Therefore, when performing *χ*^2^-independence tests, we use only the neurons present in both the current Allen Cell Type Database and the list of neurons shared by Gouwens *et al*. A total of 430 neurons matched this criterion (230 spiny and 200 aspiny).

In all datasets, the morphological information of each neuron is stored as a set of 3D coordinates with the connectivity specified in the parent samples. Complete reconstruction of neuron morphology is made by connecting data points based on their parent-child relationship. Reconstructed neuronal morphology is analyzed to identify branching points, where more than one child points identify as a parent. Tips of the neuron were defined if no other child points were present. In this study, a branch is defined as a set of points between two branching points or between a branching point and a tip.

### *F* (*q*)-based clustering of neurons

After calculating the *F* (*q*) for each neuron, we cluster the set of neurons using a hierarchical clustering algorithm. We define the ‘distance’ between the *F* (*q*)s of *i*-th and *j*-th neurons by using the Euclidian distance (L2 norm),

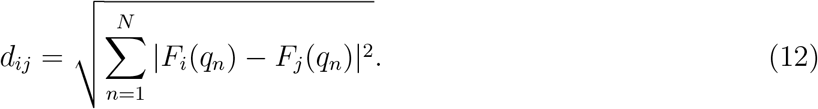

Since our interest largely lies in the intermediate *q* regime where the features of branched polymers are observed, *d*_*ij*_’s are calculated over the range of 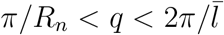. Calculation of *d*_*ij*_s over all *I* and *j* pairs gives rise to a square distance matrix, to which the hierarchical clustering algorithm is applied. For the Allen Cell type dataset, we follow the same procedure as Gouwens *et al*., who separated spiny and aspiny/sparsely spiny neurons before clustering [11]. Hierarchical clustering was carried out separately on spiny and aspiny neurons, and the clusters were collected based on the dynamic hybrid cut tree method [89]. The minimal cluster size of 4 was imposed for the mouse V1 in Gouwens *et al*. [11] and smaller clusters are automatically merged to a neighboring cluster. For the neurons in *C. elegans* and *Drosophila* olfactory system, we applied the same criterion of clustering as that of the mouse V1 neurons for consistency but did not impose the minimal cluster size restriction so that if necessary we can sample small clusters as well.

### Pearson’s *χ*^2^-test of independence

In this study, Pearson’s *χ*^2^-test is used to evaluate if there is any statistically significant relationship between the two clustering outputs, one from our *F* (*q*)-based analysis and another from Gouwens *et al* ‘s [11], which are summarized in the contingency tables for the cases of spiny and aspiny neurons in mouse V1 (see Supplementary Tables S1 and S2). For the null hypothesis, it is assumed that the clustering outputs obtained from the two different methods are statistically independent of each other, such that knowing a clustering output from one method does not help us predict the output of another method. For the statistical evaluation of independence between two clustering outputs, we calculate the *χ*^2^ value of the data in the contingency tables. If the two clustering outputs were not random, a large value of *χ*^2^ that greatly deviates from the expected *χ*^2^ distribution would be obtained. Specifically, the *χ*^2^ value is calculated using

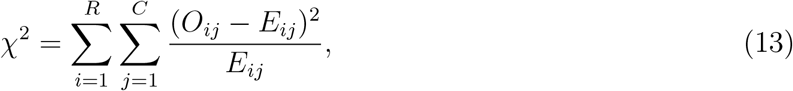

where *R* and *C* are the numbers of rows (Gouwens *et al*.’s clustering outputs) and columns (*F* (*q*)based clustering outputs) of the contingency table, respectively, and *O*_*ij*_ and *E*_*ij*_ are the observed and expected frequencies of the neurons in the *i*-th row and the *j*-th column of the table. The expected frequency *E*_*ij*_(= *Np*_*i*·_*p*_·*j*_) would be obtained under the assumption that the null hypothesis is correct (two clustering labels are independent), where 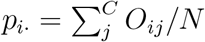 and 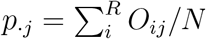 with *N* being the total number of neurons. Once the *χ*^2^ value is calculated with Eq. 13, the probability of obtaining a *χ*^2^ value larger than what is expected from the null hypothesis, namely, the p-value is calculated from

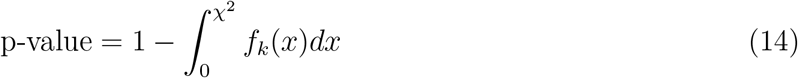

where 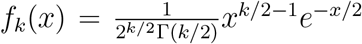 is the *χ*^2^-distribution with *k*[= (*R* − 1)(*C* − 1)] denoting the degree of freedom of the contingency table. If the p-value is smaller than a chosen significance level, the null hypothesis is rejected, and one can claim that there is a statistically significant association between the two clustering outputs.

## Declarations

### Authors’ contributions

K.C., W.K.K. and C.H. wrote the main manuscript text and K.C. prepared figures. All authors reviewed the manuscript.

### Ethics approval

Not applicable.

### Consent to participate

Not applicable.

### Data availability statement

Python scripts used for this manuscript and the complete neuron morphologies in each cluster for the *C. elegans* nervous system, *Drosophila* PNs, and mouse V1 neuron are available at https://github.com/kirichoi/FqClustering.

## Acknowledgments

We thank the Center for Advanced Computation in KIAS for providing computing resources.

