## Supplementary Text and Tables for "Polymer physics-based classification of neurons"

### Alternative expressions for form factor

The $F(q)$ in Eq. 1 corresponds to the Fourier transform of the density-density correlation function of monomers at two positions. By introducing the density field $\rho(\mathbf{r}_\alpha) = \sum_{i=1}^{N} \delta(\mathbf{r}_\alpha - \mathbf{r}_i)$ ($\alpha = 1,\, 2$), one can write Eq. 1 as

$$
\begin{aligned}
F(q) &= \frac{1}{N} \int \int \langle \rho(\mathbf{r}_1)\rho(\mathbf{r}_2) \rangle e^{-i\mathbf{q}\cdot(\mathbf{r}_1 - \mathbf{r}_2)} d\mathbf{r}_1 d\mathbf{r}_2 \\
&= \frac{1}{N} \int \int \langle \rho(\mathbf{r}_{12})\rho(\mathbf{r}_2) \rangle e^{-i\mathbf{q}\cdot\mathbf{r}_{12}} d\mathbf{r}_{12} d\mathbf{r}_2 \\
&= \int p(\mathbf{r}) e^{-i\mathbf{q}\cdot\mathbf{r}} d\mathbf{r}. \qquad (1)
\end{aligned}
$$

Here, the term $\langle \rho(\mathbf{r}_{12})\rho(\mathbf{r}_2) \rangle$ is the joint probability that one monomer is located at $\mathbf{r}_{12} (\equiv \mathbf{r}_1 - \mathbf{r}_2)$ relative to another monomer at position $\mathbf{r}_2$, which is equivalent to the conditional probability of the monomer at $\mathbf{r}$ given that another monomer is at the origin ($\mathbf{r}_0$) multiplied by the mean local monomer density, i.e. $\frac{1}{N} \int \int \langle \rho(\mathbf{r}_{12})\rho(\mathbf{r}_2) \rangle e^{-i\mathbf{q}\cdot\mathbf{r}_{12}} d\mathbf{r}_{12} d\mathbf{r}_2 = \frac{1}{N} \int \langle \rho(\mathbf{r}|\mathbf{r}_0) \rangle e^{-i\mathbf{q}\cdot\mathbf{r}} d\mathbf{r} \int \rho(\mathbf{r}_0) d\mathbf{r}_0 = \int p(\mathbf{r}) e^{-i\mathbf{q}\cdot\mathbf{r}} d\mathbf{r}$ with $\frac{1}{N} \int \rho(\mathbf{r}_0) d\mathbf{r}_0 = 1$ and $p(\mathbf{r}) \equiv \rho(\mathbf{r}|\mathbf{r}_0)$. Hence, the form factor is equivalent to the Fourier transform of pair distance distribution $p(\mathbf{r})$ as in Eq. 2.

Table 1: Contingency table of the spiny neurons in mouse V1.

| | $C_1$ | $C_2$ | $C_3$ | $C_4$ | $C_5$ | $C_6$ | $C_7$ | $C_8$ | $C_9$ | $C_{10}$ | $C_{11}$ | Total |
|---|---|---|---|---|---|---|---|---|---|---|---|---|
| Spiny_1 | 0 | 1 | 0 | 0 | 0 | 0 | 2 | 0 | 1 | 4 | 0 | 8 |
| Spiny_2 | 1 | 4 | 0 | 1 | 0 | 1 | 0 | 0 | 0 | 1 | 0 | 8 |
| Spiny_3 | 8 | 2 | 2 | 0 | 2 | 0 | 1 | 0 | 0 | 0 | 0 | 15 |
| Spiny_4 | 2 | 1 | 8 | 0 | 2 | 0 | 4 | 0 | 0 | 1 | 0 | 18 |
| Spiny_5 | 0 | 1 | 8 | 0 | 9 | 0 | 2 | 0 | 0 | 0 | 0 | 20 |
| Spiny_6 | 0 | 0 | 1 | 1 | 3 | 1 | 0 | 0 | 0 | 0 | 0 | 6 |
| Spiny_7 | 9 | 0 | 4 | 1 | 2 | 0 | 1 | 0 | 1 | 1 | 0 | 19 |
| Spiny_8 | 7 | 3 | 3 | 0 | 1 | 0 | 0 | 0 | 0 | 0 | 0 | 14 |
| Spiny_9 | 2 | 8 | 1 | 0 | 0 | 0 | 2 | 3 | 0 | 0 | 0 | 16 |
| Spiny_10 | 1 | 0 | 0 | 0 | 0 | 0 | 0 | 0 | 0 | 0 | 0 | 1 |
| Spiny_11 | 0 | 1 | 0 | 1 | 0 | 0 | 1 | 2 | 2 | 0 | 1 | 8 |
| Spiny_12 | 2 | 7 | 0 | 4 | 0 | 0 | 0 | 2 | 2 | 0 | 0 | 17 |
| Spiny_13 | 1 | 1 | 0 | 0 | 0 | 0 | 0 | 3 | 1 | 0 | 0 | 6 |
| Spiny_14 | 0 | 1 | 0 | 3 | 0 | 1 | 0 | 4 | 6 | 2 | 0 | 17 |
| Spiny_15 | 3 | 0 | 1 | 1 | 1 | 0 | 2 | 0 | 0 | 0 | 1 | 9 |
| Spiny_16 | 0 | 1 | 0 | 4 | 0 | 1 | 0 | 0 | 0 | 0 | 0 | 6 |
| Spiny_17 | 5 | 3 | 0 | 3 | 0 | 5 | 0 | 0 | 0 | 0 | 0 | 16 |
| Spiny_18 | 2 | 4 | 0 | 4 | 0 | 6 | 0 | 0 | 0 | 0 | 0 | 16 |
| Spiny_19 | 0 | 0 | 0 | 5 | 0 | 1 | 0 | 0 | 0 | 0 | 4 | 10 |
| Total | 43 | 38 | 28 | 28 | 20 | 16 | 15 | 14 | 13 | 9 | 6 | 230 |

Table 2: Contingency table of the aspiny neurons in mouse V1.

| | $C_1$ | $C_2$ | $C_3$ | $C_4$ | $C_5$ | $C_6$ | $C_7$ | $C_8$ | Total |
|---|---|---|---|---|---|---|---|---|---|
| Aspiny_1 | 7 | 1 | 0 | 0 | 0 | 0 | 2 | 0 | 10 |
| Aspiny_2 | 7 | 0 | 0 | 0 | 0 | 0 | 3 | 0 | 10 |
| Aspiny_3 | 5 | 0 | 0 | 0 | 3 | 0 | 0 | 0 | 8 |
| Aspiny_4 | 12 | 2 | 0 | 0 | 1 | 0 | 4 | 0 | 19 |
| Aspiny_5 | 1 | 7 | 0 | 0 | 0 | 2 | 0 | 0 | 10 |
| Aspiny_6 | 0 | 1 | 0 | 0 | 0 | 3 | 0 | 0 | 4 |
| Aspiny_7 | 2 | 8 | 0 | 0 | 0 | 2 | 0 | 0 | 12 |
| Aspiny_8 | 8 | 9 | 0 | 0 | 0 | 2 | 4 | 0 | 23 |
| Aspiny_9 | 1 | 2 | 0 | 0 | 0 | 4 | 0 | 0 | 7 |
| Aspiny_10 | 3 | 2 | 0 | 0 | 0 | 1 | 1 | 0 | 7 |
| Aspiny_11 | 3 | 1 | 0 | 0 | 3 | 2 | 0 | 0 | 9 |
| Aspiny_12 | 6 | 0 | 0 | 0 | 0 | 0 | 5 | 0 | 11 |
| Aspiny_13 | 1 | 0 | 2 | 1 | 1 | 3 | 0 | 0 | 8 |
| Aspiny_14 | 0 | 0 | 1 | 5 | 6 | 0 | 0 | 0 | 12 |
| Aspiny_15 | 0 | 0 | 0 | 2 | 3 | 1 | 0 | 0 | 6 |
| Aspiny_16 | 3 | 0 | 2 | 0 | 8 | 1 | 0 | 0 | 14 |
| Aspiny_17 | 2 | 2 | 0 | 0 | 2 | 1 | 1 | 0 | 8 |
| Aspiny_18 | 1 | 6 | 0 | 0 | 0 | 6 | 0 | 0 | 13 |
| Aspiny_19 | 4 | 2 | 0 | 0 | 2 | 0 | 1 | 0 | 9 |
| Total | 66 | 43 | 5 | 8 | 29 | 28 | 21 | 0 | 200 |
